# Penalized reduced rank regression for multi-outcome survival data supports a common metabolic risk score for age-related diseases

**DOI:** 10.1101/2024.11.11.622920

**Authors:** Marije H. Sluiskes, Hein Putter, Marian Beekman, Jelle J. Goeman, Mar Rodríguez-Girondo

## Abstract

The increasing availability of multi-outcome data in health research presents new opportunities for understanding complex health processes, such as ageing. Ageing is a multifaceted process, encompassing both lifespan and healthspan, as well as the onset of age-related diseases. To model this complexity, we propose the penalized reduced rank regression model for multi-outcome survival data (penalized survRRR), which identifies shared latent factors driving multiple outcomes. The model imposes a rank constraint on the coefficient matrix to capture underlying mechanisms of ageing, while accommodating high-dimensional and correlated predictors and outcomes by introducing penalization. We discuss the statistical properties of this doubly-regularized approach and show how the optimal number of ranks can be estimated from the data. We apply a lasso-penalized reduced rank regression model to 78,553 participants of the UK Biobank, using over 200 metabolic variables as predictors and the onset of seven age-related diseases and mortality as the outcomes of interest. Our results indicate that a rank 1 model provides the best fit to the data, resulting in a single metabolite-based score of age-related disease susceptibility. This highlights the potential of the penalized survRRR model to provide new insights into the nature of the relationship between metabolomics and age-related diseases.

## 1 Introduction

Multi-outcome data is becoming increasingly pervasive within health research. Diverse data sources such as electronic health records, genomic measurements and data from wearable devices are increasingly linked in large databases, capturing a wide array of health-related variables. As a result, it has become much easier to follow the health status and disease trajectory of study participants over time.

In ageing research, the rise of large epidemiological studies with rich multi-outcome data is particularly relevant. In recent years, the field of ageing research, while developing predictors such as ‘biological age clocks,’ has primarily focused on time-to-mortality as the sole outcome of interest^1,2,3^. The appeal of this outcome is that it is unambiguously defined and recorded in most longitudinal cohort studies. In addition, statistical methods for single-outcome time-to-event data are widely available. However, ageing entails more than lifespan alone: it is a multi-faceted phenomenon, characterized by the gradual accumulation of various physiological changes and damage, which is influenced by a complex interplay of genetic, environmental and lifestyle factors^4^. Hence, when modelling the ageing process it is interesting to not only consider lifespan, but also healthspan and the pattern of age-related diseases that occurs within an individual.

Age-related diseases are disease for which chronological age is a major risk factor. The conjecture is that is that these age-related diseases are all driven by a shared underlying cause, namely the general process of ageing. This makes it relevant to model the underlying mechanisms that influences the time to onset of age-related diseases, to obtain a more comprehensive understanding of the ageing process as a whole. Methodologically, this calls for a multi-outcome survival analysis approach that accommodates multiple event types within a single analytical framework. In addition, such an analytical approach should be able to deal with high-dimensional or strongly correlated predictors, as the predictor variables considered within the field of biological ageing often belong to the ‘omics’ field, a suffix which refers to the various types of biological molecules that make up the cells of an organism. This introduces an additional layer of statistical complexity.

To this end, we introduce the penalized reduced rank regression model for survival data, or penalized survRRR model. The penalized survRRR model is a multivariate model, fitted on *P* predictors and *Q* time-to-event outcomes. By putting a constraint on the rank of the coefficient matrix **B** (of dimensions *P* × *Q*), the model assumes that there is a set of *R* shared latent factors that drives all outcomes considered. This is achieved by decomposing **B** as **AΓ**^⊤^ and constraining the dimensions of the matrices **A** to *P* × *R* and **Γ** to *Q* × *R*. The penalized survRRR model can provide insight into the latent factors that underlie the age-related diseases, thereby shedding light on the underlying mechanisms of ageing. By adding a penalty term in the estimation of both **A** and **Γ**, the inclusion of high-dimensional and/or strongly correlated predictors or outcomes is possible. The proposed method is hence doubly regularized. Although we apply the penalized survRRR model to age-related disease onset data in this paper, the penalized survRRR model can be fitted to any type of multi-outcome survival data.

A reduced rank regression model has been fitted on survival data before in a competing risks setting^5^ and in a multi-state setting^6^, and a group lasso-penalized reduced rank model has been proposed for a large-scale linear regression setting^7^. However, to the best of our knowledge a penalized version of the reduced rank model for multi-outcome survival data does not yet exist. In addition, in this paper we show that when considering a penalized survRRR model where the penalty is not rotation invariant (such as the lasso penalty), the resulting decomposition of **B** into **A** and **Γ** is unique (up to trivial transformations that do not affect the interpretation). This is not the case for the unpenalized survRRR model, where an additional constraint is needed to require a unique solution for **A** and **Γ**.

In our motivating application, we consider metabolic variables as our predictor variables of interest. Referred to as “the link between genotype and phenotype”^8^, metabolite measurements are generally easy to obtain (requiring only a blood draw) and are therefore considered attractive biomarkers^9^.

Several other studies have recently attempted to assess the relation between metabolomic predictors and time-to-disease-onset outcomes, but none of them address our aim of directly modelling the sharent latent process underlying all age-related diseases and mortality. van den Berg et al.^10^ considered time to two age-related diseases as the outcome of interest. This requires an a priori assumption that the included diseases are comparable and in some sense interchangeable, such that any combination of two age-related diseases can be classified under the same name, i.e. ‘multimorbidity’. The penalized survRRR model does not require such an assumption beforehand. Pietzner et al.^11^ fit a separate model for each age-related disease and investigated which predictors were associated with more than one age-related disease. This approach provides valuable insights into the number of predictors that are associated with multiple age-related diseases, but it is not the most efficient (it requires a multiple-testing correction) and is focused on inference only. Buergel et al.^12^ trained a multitask neural network, consisting of a shared network for all diseases and disease-specific head networks. The appeal of their model is that it can deal with nonlinear associations, which the penalized survRRR model cannot. The downside is the lack of interpretability, as it focuses on prediction only. The neural network also cannot provide insight into the number of underlying shared latent factors. In contrast, the penalized survRRR model can be used for both inference and prediction, as it aims to model the underlying mechanism.

The subsequent sections of this paper are structured to provide a comprehensive understanding of the penalized survRRR model. The next section discusses the characteristics, interpretation and assumptions of a general reduced rank regression model and provides details on the non-penalized reduced rank regression model for survival data and the penalized reduced rank model in a linear regression setting. This is followed by an in-depth discussion on the penalized survRRR model, including considerations on the penalty term, the uniqueness of the lasso-penalized model solution and computational aspects. In Section 3 the penalized survRRR model is applied to age-related morbidity event data from the UK Biobank (UKB), utilizing blood metabolites as our predictors of interest. We conclude with a discussion.

## 2 Methods

### 2.1 Statistical background

#### 2.1.1 Reduced rank regression

The reduced rank regression (RRR) model was originally proposed by^13^ in the context of multivariate linear regression. When fitting a regression model on *Q* different outcomes and *P* different predictors, the resulting coefficient matrix **B** will be of dimensions *P* × *Q*. The *Q* columns contain the regression coefficients for each outcome. These regression coefficients are the same as when a separate univariate model would have been fitted for each outcome.

In contrast, when fitting a reduced rank regression model, coefficient matrix **B** is constrained to be of maximal rank *R*, where *R* ≤ min(*P, Q*). This rank constraint can be achieved by decomposing **B** as **AΓ**^⊤^, where **A** ∈ ℝ^*P* ×*R*^ and **Γ** ∈ ℝ^*Q*×*R*^. This is equivalent to the assumption that there is a set of *R* latent factors that drive all *Q* outcomes considered and is in fact the same as a multilayer perceptron with a single hidden layer and a linear activation function^14^.

The reduced rank model can conveniently be visualized in such a way that its network-like structure immediately becomes apparent. In Figure 1 a reduced rank model of rank *R* is shown, with *P* predictors, *x*_1_, .., *x*_*P*_ and *Q* outcomes, *y*_1_, …, *y*_*Q*_. When decomposing coefficient matrix **B** as **AΓ**^⊤^, **A** contains the effects of the *P* predictors on the *R* latent factors and **Γ** contains the effects of the *R* latent factors on the *Q* outcomes.

**Figure 1.**
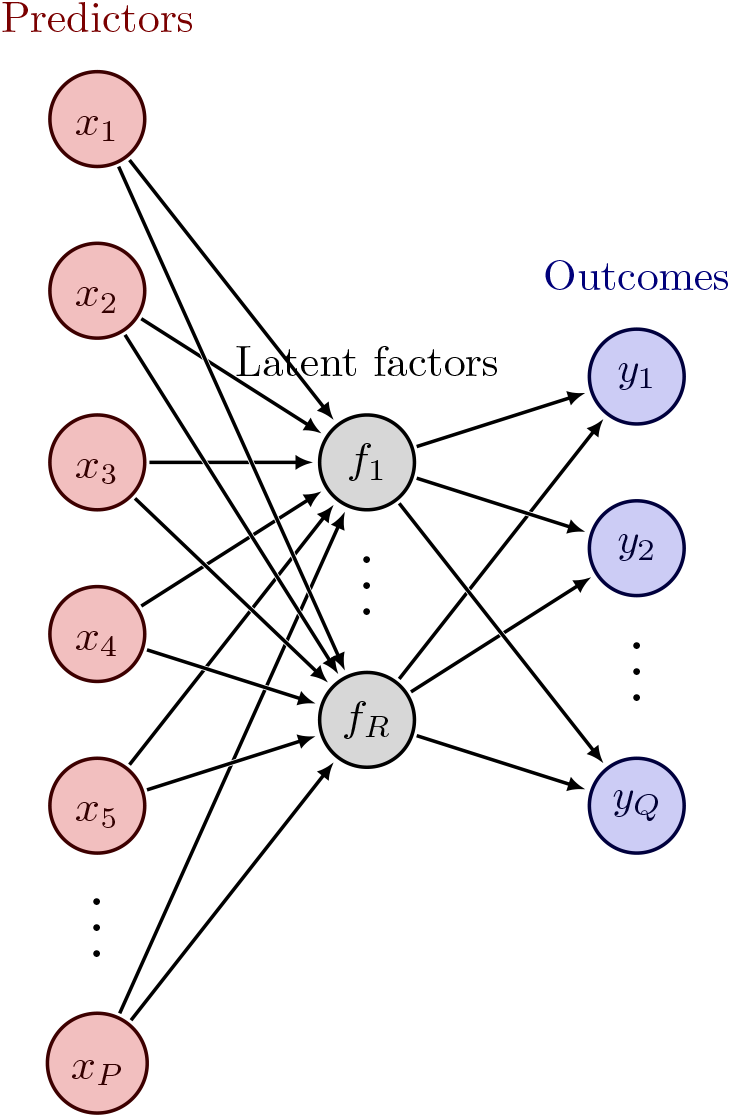
Visualization of a reduced rank model of rank *R*, with *P* predictors and *Q* outcomes.

There are several appealing aspects to the reduced rank model. Firstly, the number of coefficients that have to be estimated is reduced: whereas a full-rank model has *P* × *Q* regression coefficients, a rank *R* model only has *R*(*P* + *Q* − *R*)^5^. Furthermore, the reduced rank regression model facilitates interpretation of the parameter estimates: each latent factor might have a distinct (biological) interpretation.

#### 2.1.2 RRR for survival data

Fiocco et al.^5^ extended the reduced rank regression model of Anderson^13^ to a survival analysis context, considering time-to-event outcomes. They considered a competing risks setting, i.e. as soon as a subject in the data experiences an event for one of the *Q* different outcomes, they also disappear from the risk sets of all other outcomes.

Similar to the linear regression setting, in a time-to-event setting it is also convenient to decompose coefficient matrix **B** in a matrix **A** and a matrix **Γ** to achieve the rank constraint that rank(**B**) ≤ min(*P, Q*). For a single Cox model, the hazard for outcome *q* = 1, …, *Q* for an individual with vector of covariates *X* = (*x*_1_, …, *x*_*P*_) is

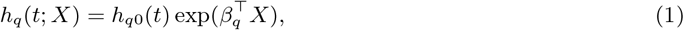

where *h*_*q*0_(*t*) is the baseline hazard for event type *q* and *β*_*q*_ is a vector of length *P*, containing the regression coefficients for this event. The *P* × *Q* regression coefficients of the *Q* Cox models can be represented in a matrix **B**, where *β*_*q*_ is the *q*^th^ row of **B**. When again decomposing **B** as **AΓ**^⊤^, with **A** = [*α*_1_|…|*α*_*R*_] and **Γ** = [*γ*_1_|…|*γ*_*R*_], Equation (1) can be rewritten as

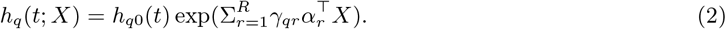

Note that *γ*_*qr*_ is a scalar, as the model is stratified by the number of outcomes *Q*, while *α*_*r*_ is *P* -length vector, which is shared across all outcomes.

Now the task is to estimate **A** and **Γ** (and hence **B**). Fiocco et al.^5^ used an iterative procedure where the likelihood is alternately optimized with respect to **A** and **Γ**: first **Γ** is taken as fixed and the maximum likelihood solution for **A** is found (using *γ*_*qr*_ *X* as covariates), then this newly found solution for **A** is taken as fixed and the maximum likelihood solution for **Γ** is found (using *α*_*r*_^⊤^*X* as covariates). As remarked by Fiocco et al.^5^, this idea is known under different names, such as the NIPALS algorithm^15^, the criss-cross method^16^ and the partitioned algorithm^17^.

Even though the coefficient matrix **B** is generally uniquely defined, the decomposition **B** = **AΓ**^⊤^ is not, since for any *R* × *R* invertible matrix 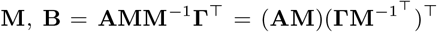.Hence, additional constraints on the decomposition are needed to obtain a set of unique parameter values for **A** and **Γ**, e.g. by using the singular value decomposition. See Yee and Hastie^18^ for an overview of other possible uniqueness constraints.

When using the RRR model to analyse time-to-event data, an additional advantage of this model is that it can ‘borrow strength’ when the data contains outcomes with few events, in particular when *R* is small. Few events will lead to imprecise estimates, but when fitting a reduced rank model, the events of all outcomes are used to obtain an estimate for **A**. Each row in **Γ** is estimated using the events for outcome *q* only.

#### 2.1.3 Penalized RRR models

The rank constraint on RRR models is a form of regularization. However, in certain cases the coefficient matrix **B** may still become unidentifiable, e.g. when the number of predictors *P* is nearing or exceeding the number of rows in the data *n*, or difficult to estimate, e.g. when predictors are strongly correlated. A popular approach in such cases is to fit a penalized model, in which a penalty term is added to the original objective function. In that case the RRR model becomes doubly regularized.

A sparse reduced rank linear regression model for large-scale ultrahigh-dimensional problems (SRRR) has recently been introduced by Qian et al.^7^. They fitted a group-lasso penalized RRR model on data from the UK Biobank, using genetic data (SNPs) as predictors and phenotypic data (35 biomarkers) as outcomes. The chosen group-lasso approach treats the effect of each predictor on all outcomes as a distinct group: i.e. either the predictor has an effect on all outcomes or it has an effect on none, to achieve homogeneous sparsity across outcomes.

Other choices for the penalty term can be more suitable in other application contexts. The most popular penalties are the ridge penalty^19^, which penalizes the squared values of the coefficients and shrinks coefficients towards zero, the lasso penalty^20^, which penalizes the absolute values of the coefficients, and the elastic net penalty^21^, which can be considered a compromise between the ridge and lasso penalty.

### 2.2 Penalized RRR models for survival data

The previous section discussed relevant models and developments within the context of reduced rank regression. This section introduces our new model: a penalized reduced rank regression model for survival data, which we have named penalized survRRR.

#### 2.2.1 Model characteristics

In the penalized survRRR model, a penalty term is placed on both matrix **A** and matrix **Γ**, where **B** = **AΓ**^⊤^. The corresponding penalization parameters are denoted by λ_**A**_ and λ_**Γ**_. The penalized log partial likelihood of the model can hence be written as follows:

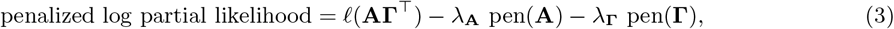

where *ℓ* denotes the log partial likelihood and pen denotes some penalty. Note that a stratified (penalized) Cox model is fit, which means that the total log partial likelihood *ℓ* is the sum of the usual Cox model log partial likelihoods of the individual outcomes *q*.

When fitting the model, the aim is to find **A** and **Γ** such that the penalized log partial likelihood in Equation (3) is maximized, or alternatively, that the penalized deviance (−2 times the negative log partial likelihood) is minimized. The λs are free tuning parameters: the larger their value, the more the coefficients of **A** and **Γ** are shrunk towards zero. The model performance suffers if λ is either too small or too large.

Once the model has been fit and the optimal **A** and **Γ** have been found, there exist matrices **A**^∗^ and **Γ**^∗^ that result in the same **B**: as given in Section 2.1.2, for any invertible matrix 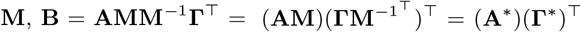. Hence, the likelihood-part of Equation (3), *ℓ*(**AΓ**^⊤^), does not depend on whether **A** and **Γ** or **A**^∗^ and **Γ**^∗^ are chosen.

Whether the two penalty terms in Equation 3 change when choosing **A**^∗^ and **Γ**^∗^ instead of **A** and **Γ** depends on the penalty. If the constraint region associated with the penalty is invariant to rotation, the non-identifiability issue inherent to the non-penalized version of the (surv)RRR model remains, i.e. if we take **M** to be a rotation matrix, λ_**A**_pen(**A**) + λ_**Γ**_pen(**Γ**) = λ_**A**_pen(**A**^∗^) + λ_**Γ**_pen(**Γ**^**∗**^). Hence, any rotation of **A** and **Γ**, resulting in a new solution with matrices **A**^∗^ and **Γ**^∗^, has the same penalized likelihood value. However, if the constraint region associated with the penalty is not invariant to rotation, rotating **A** and **Γ** results in a solution with a lower penalized likelihood. The ridge penalty is invariant to rotation; the lasso penalty is not. This implies that there are multiple solutions with the same penalized likelihood when using a ridge penalty, while there is only one solution for the lasso penalty (up to a trivial permutation of the columns of **A** and **Γ** or a multiplication of **A** by *c* and **Γ** by 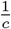, where *c* is any constant).

An intuitive illustration of the differences between the ridge and lasso penalty is provided in Figure 2, which visualizes their respective constraint regions. As mentioned, a rotation of matrices **A** and **Γ** by **M** does not move the location of the maximum likelihood solution 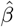, because the (log partial) likelihood does not change. However, the coordinate system does change due to the rotation: the origin remains at the same location, but the axes rotate and may shrink or stretch. The constraint region will rotate with the axes. When pen is a ridge-type penalty this does not matter, since the constraint region is a circle (or more generally an *n*-sphere). However, when pen is a lasso-type penalty it does matter: the constraint region now is a square (or more generally a cross-polytope). If this region is rotated, the penalization parameter λ must be decreased (the area must become bigger) to stay on the same red contour line and hence result in the same penalized (log partial) likelihood value.

**Figure 2.**
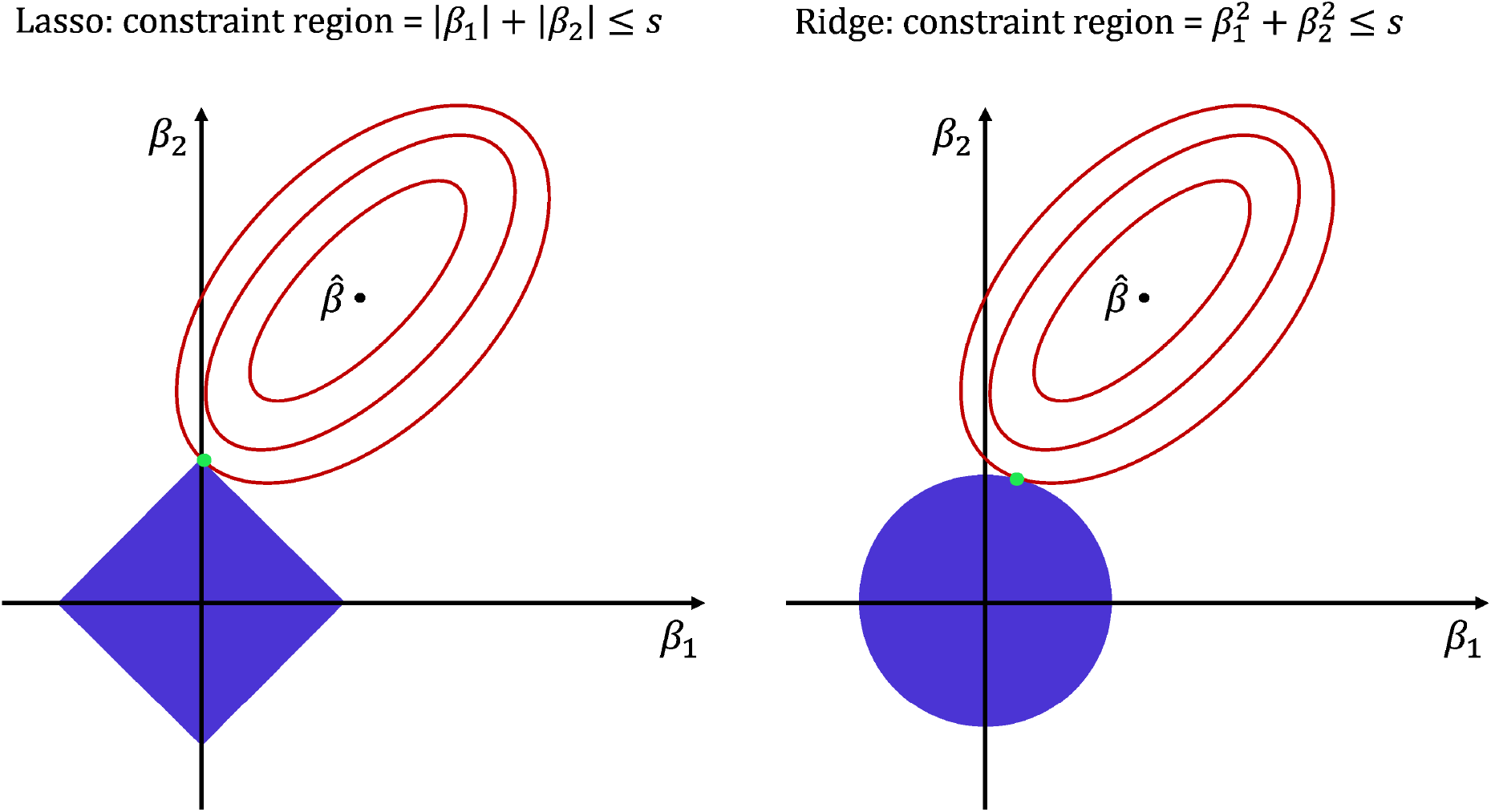
Visualization of the lasso (left panel) and ridge (right panel) penalties in terms of the constraint region, defined by *s*. The blue areas represent the constraint regions: a square for lasso, a circle for ridge. The red ellipses represent the contours of the log (partial) likelihood as it moves away from the optimal solution 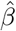. The green dot represents the lasso or ridge coefficients.

Instead of being rotated, **A** and **Γ** can also be multiplied by some constant. Although only the ridge-penalized solution is invariant to rotation, both ridge- and lasso-penalized solutions are invariant to multiplication by a constant when λ_**A**_ and λ_**Γ**_ are free parameters. This can be seen when taking **M** to be equal to some constant *c* times the identity matrix **I**. Then 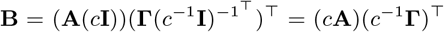.Define **A**^∗^ = *c***A** and 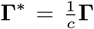, such that **B** = **A**^∗^**Γ**^∗⊤^. The penalized likelihood remains the same if 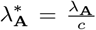 and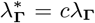.. This holds for any value of *c*, and hence, there exist many different solutions which are all scaled versions of one another. However, obtaining a unique solution is straightforward: *c* can always be chosen such that λ_**A**_ = λ_**Γ**_ without loss of generality (i.e. keeping the same penalized likelihood value). Taking λ_**A**_ = λ_**Γ**_ has as an additional advantage that it greatly simplifies fitting the (lasso- or ridge-)penalized survRRR model: to find the optimal level of penalization, the model now only needs to be fit over a sequence of λ-values instead of a two-dimensional matrix with combinations of values for λ_**A**_ and λ_**Γ**_.

Taken together, the properties of the lasso-penalized survRRR model imply that for a given value of λ (= λ_**A**_ = λ_**Γ**_), the resulting **A** and **Γ** are unique (up to a permutation of the columns): no other solution exists with the same penalized likelihood value. This also holds for any other penalty not invariant to rotation. The ridge-penalized survRRR model, which is invariant to rotation, requires an additional constraint to obtain a unique decomposition of coefficient matrix **B**, similar to the non-penalized survRRR model.

#### 2.2.2 Estimation procedure

The estimation procedure for fitting the penalized survRRR model, for a given number of ranks *R* and a given value for the penalization parameter λ, is described by Algorithm 1. In short, the penalized likelihood is alternately optimized with respect to **A** (line 3) and **Γ** (line 4) until the model has converged. It is hence very similar to that described by^5^ for the unpenalized survRRR model, although here there is an additional penalization parameter λ. This parameter should be carefully tuned. To determine the optimal value for λ, the data set should be split in a training and a test part. The model should be fit on the training data only, for a sequence of λs. The optimal value for λ can be determined by checking which value for λ resulted in the best fit on the test set.

To evaluate the fit on the test set, we again consider the log partial likelihood *ℓ*. We use the method suggested by Verweij and van Houwelingen^22^ to avoid the problem of a predictive (partial) likelihood of zero when a subject in the test set experiences an event at another time point than the subjects in the training set. This method entails evaluating the log partial likelihood on the training set and the full dataset (train and test combined) separately, and subtracting the former from the latter. We scale by the number of observations in the test set, *n*^test^, for convenience and multiply by a factor −2 to obtain the deviance, in line with the objective function used in Algorithm 1. Hence, the objective function to evaluate the fit of a penalized survRRR model on the test set is given by:

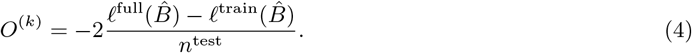

#### 2.2.3 Number of ranks

The section above described the model-fitting procedure for a given number of ranks *R*. In practice, this procedure will have to be repeated for different values of *R*. How to choose the optimal number of ranks, given the data, then follows naturally: going through the entire model-fitting procedure for a given *R* will result in an optimal 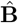. This is the coefficient matrix of the model that resulted in the lowest value of Equation (4) in the test set, associated with a certain value for λ. This procedure can be repeated for different *R*, after which the performance of the resulting models on the test set can be directly compared. Note that a test set is once again essential here: deciding on the optimal number of ranks based on the same data that the model was fitted on will result in overfitting, as a higher-rank solution will always be preferred when evaluated on the same data.

##### Algorithm 1

Estimation procedure penalized survRRR model

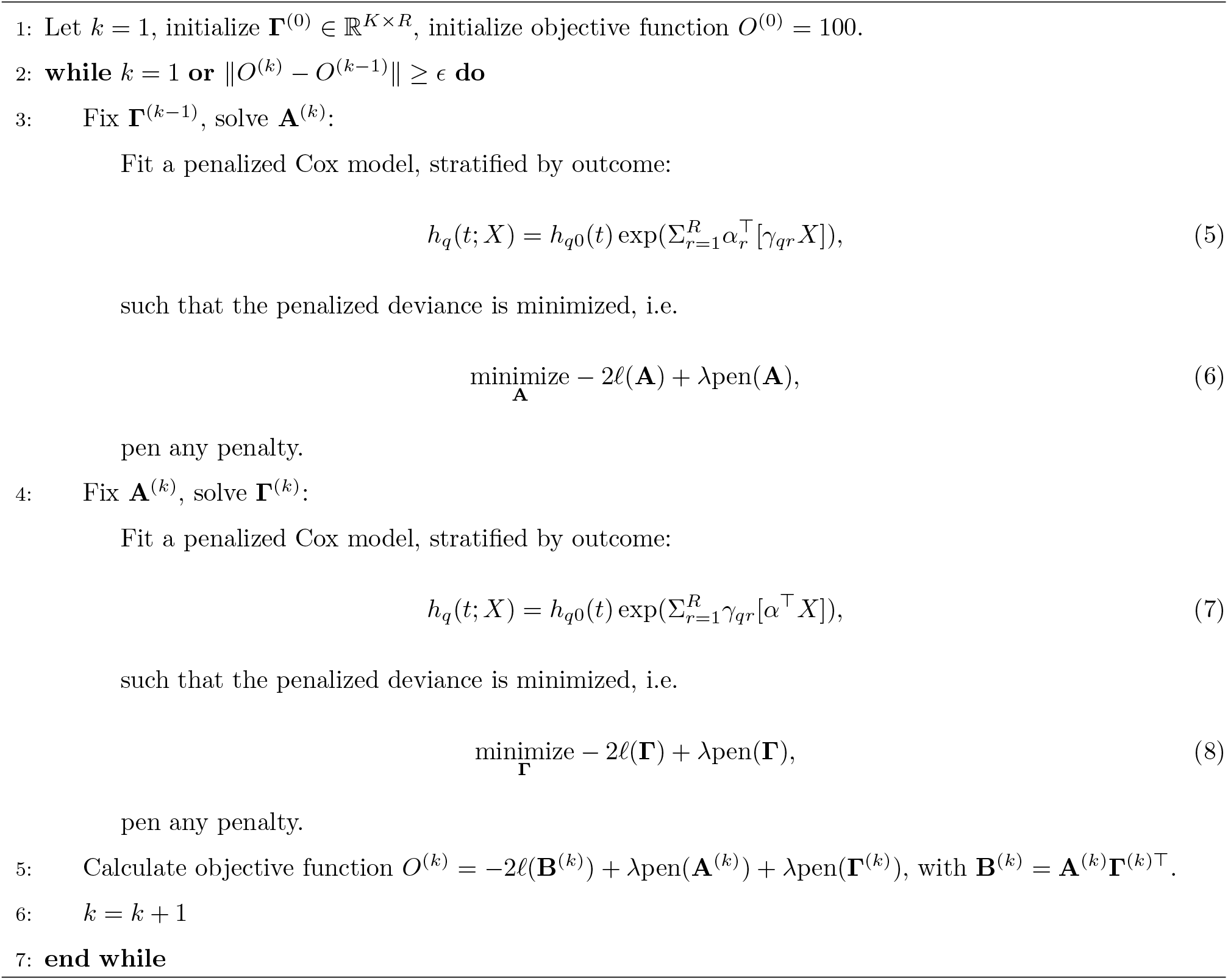

#### 2.2.4 Computational considerations

In this section we describe the most important choices that have to be made when fitting a penalized survRRR model and provide guidance on how to make them. In addition, depending on the size of the problem at hand, fitting the penalized survRRR model can prove to be somewhat of a challenge. We hence also describe the problems we encountered and the strategies we used to overcome them.

A first consideration is how to initialize the matrix **Γ** in the first iteration of Algorithm 1. The closer **Γ**^(0)^ is to the (penalized) maximum likelihood solution, the fewer iterations are needed until convergence is reached. Without any prior information, **Γ**^(0)^ can simply be initialized with values drawn from a normal or uniform distribution. It is advised to first fit the model for a relatively large value of λ. Once the model is fitted, use the final **Γ** of this converged model as the starting point for fitting the same model again with a slightly smaller λ. This is known as a so-called ‘warm start’: using the optimal value of a related (in this case: sparser, with a larger penalization parameter λ) problem to provide the initial values for a problem that is more difficult to fit (in this case: denser, with a smaller λ). Note that it is important to add a small constant to **Γ** before using it as the initial matrix **Γ**^(0)^ in the next model: for larger values of λ, one or more columns of **Γ** might become fully zero, which means that the model is not of the specified rank *R* but of some rank smaller than *R*. Adding a small constant helps to prevent that the model gets stuck in this smaller-rank solution, as we noticed that the model cannot make the ‘jump’ back to a larger-rank model itself. By adding a small constant, the model is again forced to be of the specified rank *R*.

As mentioned, the model described in Algorithm 1 should be fitted several times, namely for different values of λ, to determine the optimal amount of penalization. It is advisable to generate the values of this λ-sequence on a log-scale, such that the absolute distance between larger λ-values in the sequence is bigger than for smaller values. Finding good values for the sequence is primarily a trial-and-error process. For too small λ-values, the model will not converge. For λ-values that are too large, either **A** or **Γ** might become fully zero during the model-fitting process. If the step size between the different λ-values is too big, the model will in our experience not converge, or only very slowly, as the penalized maximum likelihood solution is too far from the initial **Γ**^(0)^.

The model can be delicate to fit. It can have issues converging, especially on larger data sets, for a larger number of ranks or for a smaller λ. In such cases it often helps to decrease the step size in between the values in the λ-sequence and/or to increase the first value. It might also help to start with a different initial **Γ**^(0)^. Nevertheless, in our experience sporadic convergence issues cannot be avoided. We therefore advise to fit several ‘runs’ of the same model-fitting procedure, slightly changing the initial conditions for each run. For example, first fit a rank *R* model with a certain λ-sequence and plot the objective function for both training (Algorithm 1, line 5) and test set (Equation (4)). Secondly, repeat this process, but slightly increase the number of λs in the sequence. Thirdly, repeat this process but initialize **Γ**^(0)^ differently, etc. Ideally, when plotting the (penalized) deviance on the training data against λ for these various runs, their lines should overlap. However, it can happen that for certain runs, the lines diverge. This implies that a certain run has not properly converged. Illustrations of these plots, including an indication of non-convergence of of of the runs, are provided in Section 3.

In theory, each iteration in Algorithm 1 should result in an improved fit. In other words, the quantity ∥*O*^(*k*)^ − *O*^(*k*−1)^∥ (line 2) should always be positive. This, however, is not always the case in practice, especially not when the model is close to convergence. We think this can be attributed to some numerical rounding issues when fitting the models in step 3 and 4. We therefore consider the model converged if the improvement becomes negative. This is a somewhat crude approach, but as several different runs are fitted for each model anyway, it will be spotted when the model had in reality not yet converged.

## 3 Application of the lasso-penalized survRRR model to UK Biobank multimorbidity data

We developed the penalized survRRR model with the specific aim to fit this model to multi-outcome age-related morbidity event data, using metabolomic variables as predictors.

Several choices can be made with respect to the penalty term. In our application, some penalty on **A** is essential, as the metabolomic variables are strongly correlated. It is therefore not possible to fit the model using the non-penalized reduced rank model for survival data. We chose to fit the model using a lasso penalty, to ensure sparsity and aid interpretability. In addition, a lasso-penalized survRRR model results in a unique solution, as explained in detail in Section 2.2.1. We also allow for a lasso penalty on the **Γ** matrix. This has as an effect that the coefficients of any age-related disease that cannot be predicted by the metabolomic variables will be set to zero.

### 3.1 Data description and preprocessing

We fitted the lasso-penalized survRRR model on data from the UK Biobank (UKB), a large-scale prospective cohort study containing data from about 500,000 participants, recruited from 22 different sites across the United Kingdom in 2006–2010. Participants were between 40 and 69 years old at time of recruitment. The UKB collects and provides access to data on the health and well-being of its participants over time. This includes extensive genotypic and phenotypic data and longitudinal follow-up for a wide range of health-related outcomes.

As predictors we used blood-based metabolic variables. In the UKB these metabolites were quantified using the high-throughput nuclear magnetic resonance (NMR) targeted metabolomics platform of Nightingale Health Ltd.^23^. Per blood plasma sample, 249 metabolic measures are provided, of which 81 are ratios. For an overview of all included metabolic measures, see the Supporting Information, Table 1. We used data from the 104,296 participants for which we had metabolic measurements at baseline available.

**Table 1:** Rows 64–67 and colums 1–3 of 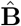 for the optimal rank 1 model (log(λ) = −11.51), given for both runs 1 and 3. The values represent the effects of the metabolic variables his, total bcaa, ile and leu (*x*_64_–*x*_67_) on diabetes, TIA and hypertension (*D*_1_–*D*_3_).

|  | (a) Run 1 |  |  |  | (b) Run 3 |  |  |
| --- | --- | --- | --- | --- | --- | --- | --- |
|  | Diabetes | TIA | Hypertension |  | Diabetes | TIA | Hypertension |
| $x_{64}$ | -0.08601 | -0.02495 | -0.03526 | $x_{64}$ | -0.08601 | -0.02496 | -0.03526 |
| $x_{65}$ | 0.00000 | 0.00000 | 0.00000 | $x_{65}$ | 0.00000 | 0.00000 | 0.00000 |
| $x_{66}$ | 0.20634 | 0.05987 | 0.08458 | $x_{66}$ | 0.20635 | 0.05987 | 0.08458 |
| $x_{67}$ | -0.30794 | -0.08934 | -0.12622 | $x_{67}$ | -0.30795 | -0.08935 | -0.12623 |

The outcomes of interest are the times to (first occurrence of) several age-related diseases and time to all-cause mortality. Information on the age at diagnosis of age-related diseases is provided by the UKB through linkage with medical and health records. We considered the same age-related diseases as van den Berg et al.^10^, based on the ICD-10 classification, but excluded breast and prostate cancer (as they are sex-specific) and cerebral infarction (as its ICD-10 code is already included in those of the transient ischemic attack ICD-10 codes). Hence, we included the following seven diseases: transient ischemic attack (TIA), angina pectoris (AP), myocardial infarction (MI), hypertension (HT), diabetes (DB), lung cancer (LC) and colon cancer (CC). We included time to all-cause mortality as the eighth outcome of interest. For an overview of the included diseases, their ICD-10 codes and corresponding UKB Field IDs, see the Supporting Information, Table 2.

**Table 2:** Matrix 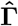 for a rank 1 model with λ equal to 1.008e-05, rounded to three decimals.

|  | DB | TIA | HP | AP | MI | LC | CC | Death |
| --- | --- | --- | --- | --- | --- | --- | --- | --- |
| $R1$ | 3.644 | 1.057 | 1.494 | 1.557 | 1.469 | 1.528 | 0.764 | 1.531 |

Metabolic variables missing in more than 1 percent of all participants were excluded, resulting in the removal of 7 out of 249 predictors. If participants had missing measurements in any of the the remaining 242 metabolic variables, they were excluded as well, removing 1,647 participants. Furthermore, participants were only included if they were 50 years or older at time of recruitment, as those below 50 years of age were considered to be too young to be diagnosed with an age-related disease during the follow-up period. This step decreased the sample size (which helped with computational feasibility) while only marginally decreasing the number of events. This left a final sample size of 78,553 participants, consisting of 36,227 men and 42,326 women. Their mean age at recruitment was 60.1 years (SD = 5.4, IQR = 56–64). Follow-up time was available until November 2021, with a median follow-up time of 13.2 years (IQR = 12.5–14.0).

Participants were at risk for a certain outcome only if they had not yet been diagnosed with this disease at time of inclusion. For example, a participant diagnosed with diabetes prior to entering the study will not be at risk for diabetes in our model, but will still be considered at risk for all outcomes that they did not yet experience (unless, naturally, the experienced outcome is death). This can be conveniently achieved by structuring the dataset in a so-called ‘long’ format, with one line per participant per outcome. See Putter et al.^24^ for a detailed explanation. After this step, the outcomes with the smallest and largest number of observed events were lung cancer (864 events) and hypertension (11,763 events), respectively. For an overview of the number of events per outcome, both before and after the age selection, see the Supporting Information, Table 3.

Given the size of the UKB data, and the large number of events for each outcome considered, we randomly split the final data set in half to create a training and a test set. Prior to the analysis, the metabolic variables were scaled and centered, separately for the training and test set.

### 3.2 Model-fitting procedure

In this section we describe how the penalized survRRR model was fit and how the optimal model was found. This provides interesting insights into the behaviour of the model and helps when fitting the model in a different context. In the next section we discuss and interpret the best-fitting model.

The optimal number of ranks is unknown. We hence started with fitting a rank 1 model. The results of fitting different runs of a rank 1 lasso-penalized survRRR model, in terms of the penalized deviance of the converged model, are provided in Figure 3. Each line represents a different run. Exact specifications of the different runs, such as λ-sequences and convergence criteria, can be found in the Supporting Information, Table 4. The left panel of Figure 3 shows the value of the objective function of a converged model on the training data (i.e. the penalized deviance, see Algorithm 1, line 5) on the y-axis, and the corresponding value for log(λ) on the x-axis. The right panel shows the performance of these same models on the test data, judged by the deviance on the y-axis (Equation (4)). For comparison, the null deviance is 1.099 on the training data and 1.254 on the test data.

**Figure 3.**
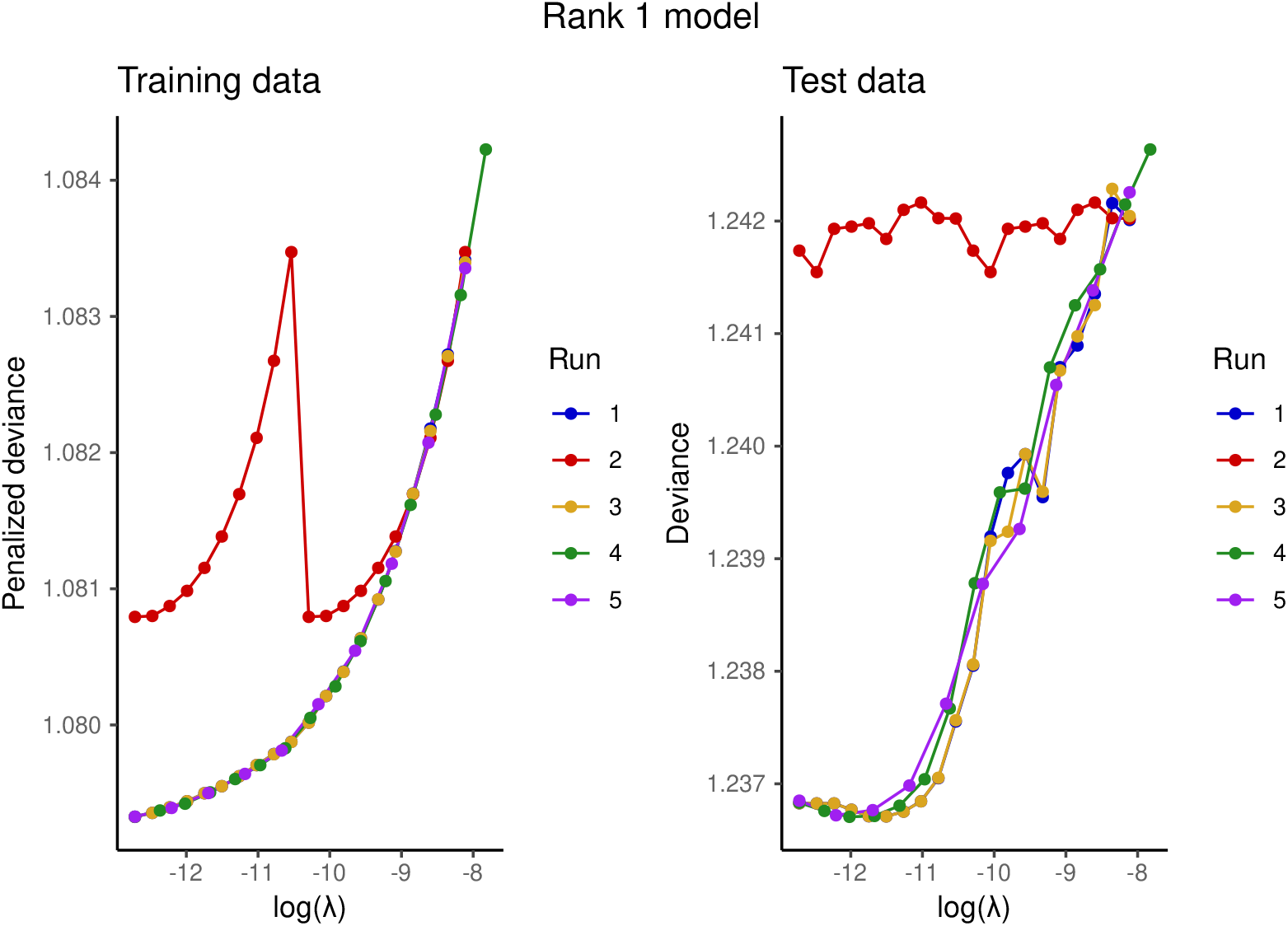
(Penalized) deviances of a converged rank 1 lasso-penalized survRRR model, for different values of λ on the x-axis. The lines represent different runs, i.e. corresponding to different initial conditions.

What becomes immediately apparent from this figure is that certain runs, which on first sight appear to be converged, can in fact deviate from the optimal path. In this example this is the case for run 2. Even small differences can induce this deviating behaviour: in fact, the only difference between run 1 and run 2 is the seed used to randomly initialize **Γ**^(0)^. This illustrates why we strongly advise to fit multiple runs, slightly changing the initial conditions in each run.

Apart from run 2, the other runs are very similar in their behaviour. Given a certain λ, their corresponding **Â** s, 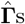 and hence 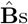 are as good as identical, as one would expect. Table 1 shows 4 rows and 3 columns of the 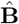 of run 1 and 3, for the value for λ corresponding with the best performance on the test data. These two runs used the same λ-sequence and can hence be directly compared. The values are close to identical, and the coefficients for some metabolic variables (*x*_65_ (total bcaa) in this case) are set to 0, due to the lasso penalty. The complete 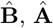 and 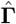 matrices are provided and discussed in the next section.

Next a rank 2 model was fitted. Figure 4 contains the results of fitting a rank 2 model lasso-penalized survRRR model. In the left panel, showing the penalized deviance on the training data, a sudden drop in penalized deviance can be seen around log(λ) = −10. This is when 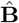 moves from a rank 1 to a rank 2 solution, i.e. to the right of this point one column of **Â** and/or 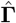 are fully 0. Moving from a rank 1 to a rank 2 solution results in a sudden improvement in fit on the training data, but in the right plot it can be seen that this results in a *lower* fit on the test data: the deviance rises sharply and only falls again as λ decreases further. The optimal rank 2 solution corresponds to a (very) small λ, implying that the final solution is almost not penalized. In fact, the penalization is so minimal that at a certain point, the model does not converge anymore on the training data. This can be spotted by the fact that the performance on the training data worsens for very small values of λ, corresponding to a not-converged model. This is when the performance on the test set also starts to increase again.

**Figure 4.**
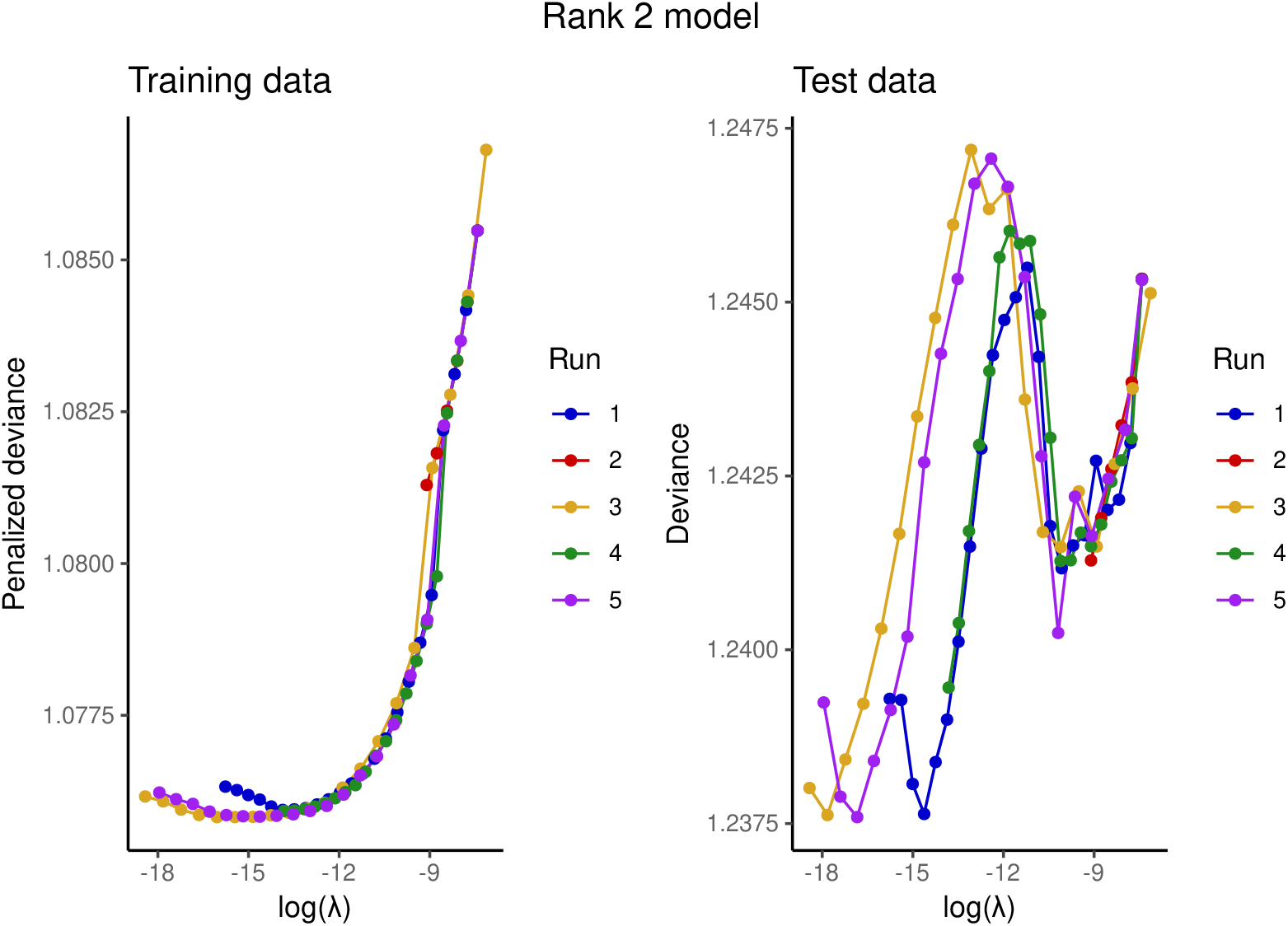
(Penalized) deviances of a converged rank 2 lasso-penalized survRRR model, for different values of λ on the x-axis. The lines represent different runs, i.e. corresponding to different initial conditions.

The converged rank 1 and rank 2 models can now be directly compared. This is necessary to determine the optimal number of ranks. The results of this comparison are shown in Figure 5. The blue lines correspond to the rank 1 solutions shown in Figure 3, the orange lines to the rank 2 solutions shown in Figure 4. The right plot illustrates that the rank 1 model is the optimal model for the data: the deviance on the test set is lowest. Furthermore, it can be seen that before the rank 2 models move to a rank 2 solution, they follow the same path as the rank 1 models, as is to be expected. From these figures it can also be concluded that it is not possible to decide on the optimal number of ranks based on the training data alone: on the training data, a higher-rank model will always result in an equal or better performance than a lower-rank model.

**Figure 5.**
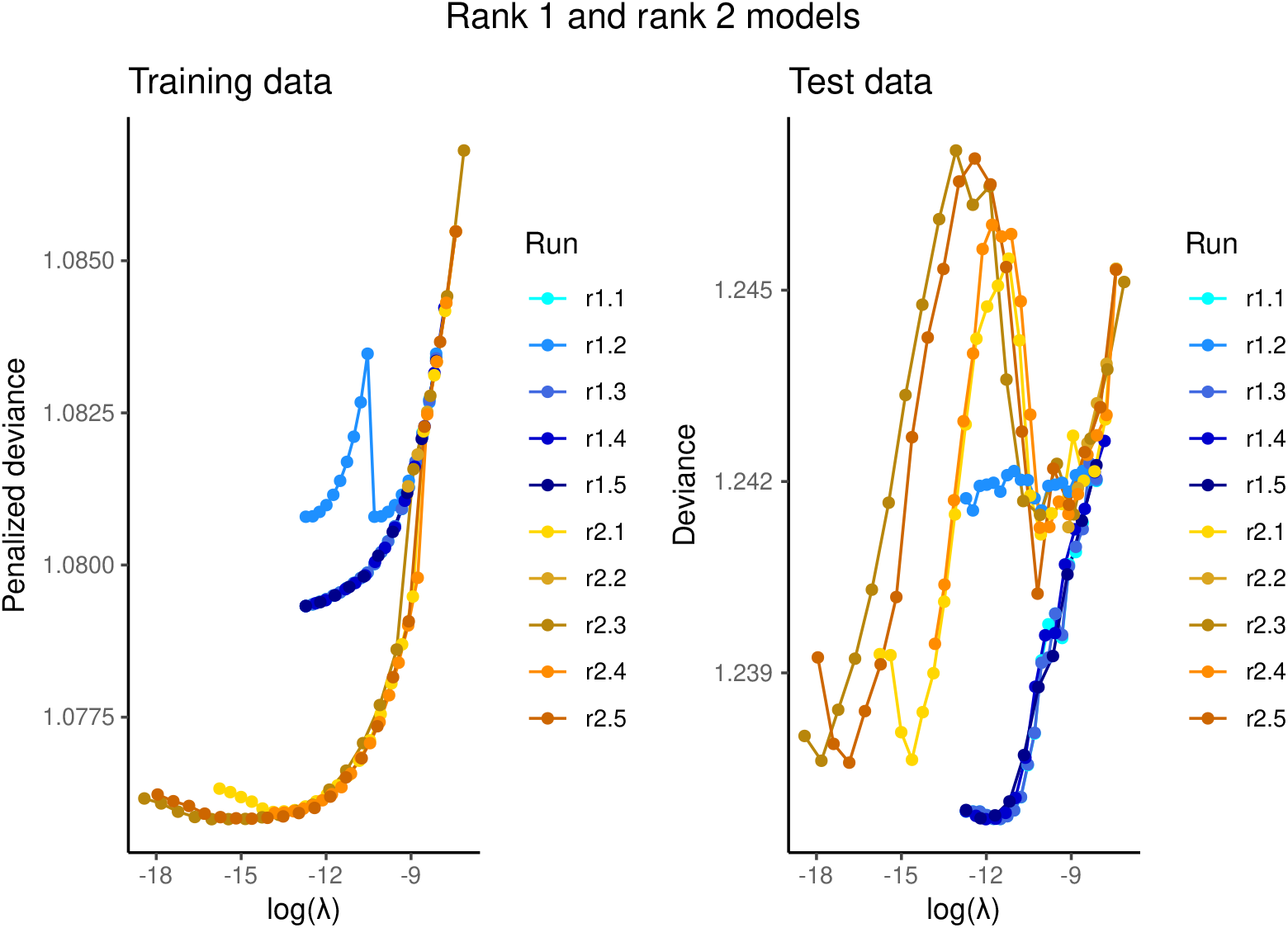
(Penalized) deviances of converged rank 1 and rank 2 lasso-penalized survRRR models, for different values of λ on the x-axis. Combined results from Figures 3 and 4.

As we found that a rank 1 model best describes the data, we have not fitted models of rank higher than 2. However, if we would have found that the rank 2 model performed better, we would have repeated the model-fitting procedure for the rank 3 model, and so forth, until the optimal number of ranks has been found. In the subsequent section, we report the results from run 3 of the rank 1 model.

### 3.3 Results

The model that best describes the data is a rank 1 model with penalization parameter λ equal to 1.008e-05. This is a small number, which is reflected by the sparsity of the corresponding coefficient matrix 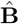 only seven of the 242 metabolites have a coefficient of 0 and none of the diseases do. Nevertheless, some penalization was essential: it was not possible to fit the unpenalized survRRR model on this data. Given its size, the full matrix 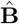 can be found in the Supporting Information, Table 5. Direct interpretation of the coefficients should be done with care, as no confidence intervals can be provided due to the penalization.

Matrix 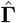, of dimensions 8 × 1, is given in Table 2. It contains the coefficients for the effect of the rank score (i.e. **Â** ^⊤^*X*) on the outcome. It can be seen that all coefficients are positive. This is in line with the conjecture that all outcomes considered are driven by the same underlying ageing process. If the rank score indeed captures the ageing process, it is to be expected that its effect on each outcome has the same direction: either all positive or all negative. In terms of coefficient size, diabetes dominates. Hypertension, angina pectoris, myocardial infarction, lung cancer and death have an associated coefficient that is about the same size. The coefficients for colon cancer and transient ischemic attack are smallest.

Directly summarizing the coefficient matrix **Â**, of dimensions 242 × 1, is not straightforward. See Table 6 of the the Supporting Information for the full matrix. A more informative approach to interpret the results is therefore to consider the rank scores of the penalized survRRR model, **Â** ^⊤^*X*, in more detail. These rank scores can for instance be compared to the linear predictors of univariate models, fitted on a single outcome. In our motivating example, this is particularly straightforward, as the optimal number of ranks is 1. There hence is only a single rank score.

As it is common practice within the ageing field to develop biological age predictors using time-to-mortality as the only outcome of interest, we compare the rank score of our optimal lasso-penalized survRRR model with the linear predictors of a lasso-penalized Cox model with time-to-mortality as the outcome and the same metabolites as predictors. The optimal value for λ was found to be 3.263e-06 for the penalized Cox model. Both models were fitted on the training data set and evaluated on the test data. Results can be found in Figure 6. As expected, there is a clear positive correlation (Pearson’s correlation coefficient = 0.79): both models were trained on the same data, with the same predictors, and the same outcome. However, the penalized survRRR model considered more than one outcome, and as can be derived from Table 2, death is not the outcome with the highest coefficient in 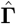. In essence, Figure 6 illustrates that the penalized survRRR model captures much of the same as a model trained on all-cause mortality alone, but not the exact same.

**Figure 6.**
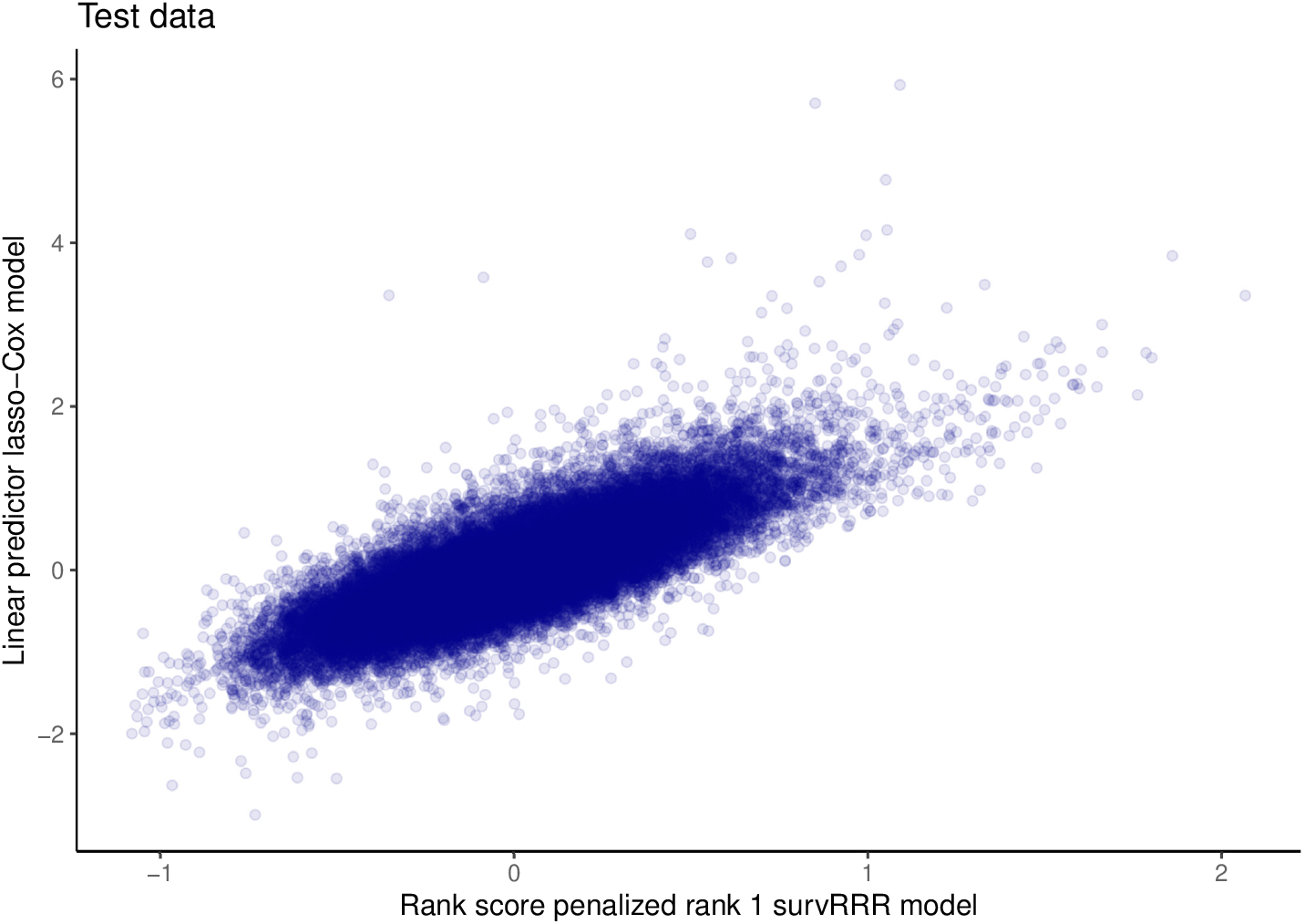
Comparison of rank scores from optimal penalized survRRR model, which is of rank 1, with the linear predictors of a lasso-penalized Cox model with mortality as the only outcome. Both models were fitted on the training data and evaluated on the test data. Opacity of the dots was set to 0.1.

Another insightful comparison is that of the penalized survRRR model rank score with a metabolic predictor fitted on other datasets than the UKB. One such predictor is the MetaboHealth score^2^. This predictor of all-cause mortality was developed starting from the same metabolic variables, using data from 44,168 individuals from 12 different cohorts. The MetaboHealth score itself is a weighted risk score based on 14 metabolic variables. A description of how the MetaboHealth score was calculated is provided in the Supplementary Information, Appendix A. Figure 7 plots the rank score of the penalized survRRR model against the MetaboHealth score in the test data. It can be seen that there is again a positive correlation (Pearson’s correlation coefficient = 0.41), albeit less strong than in Figure 6. This is not unexpected: not only is the MetaboHealth score based on time-to-mortality only, it is also based on just 14 metabolites (which keeps the score easy to interpret and apply, but it is bound to sacrifice some predictive accuracy). Most importantly, it was fitted on different data. The metabolic variable measurements are known to be 5-10% diluted in the UKB^25^, which might impact comparability, and the MetaboHealth score was constructed based on the 2014 quantification of the Nightingale Health platform, whereas the UKB contains the 2020 quantification. Although this requantification was found to have a limited effect on backward replicability of the MetaboHealth score in terms of its association with mortality^26^, it likely still affects the correlation in Figure 7. Taken together, Figure 7 paints a similar picture as Figure 6: the rank score of the optimal penalized survRRR model correlates positively with metabolic predictors of all-cause mortality, but it captures more, as it was fitted on more than one relevant outcome. Which is desirable depends on the research aim: if the goal is to predict overall mortality risk as accurately as possible, fitting a model with time-to-mortality as the only outcome of interest is the logical choice. If the goal is to capture the underlying ageing process, considering several age-related diseases in addition to death might be the more natural choice.

**Figure 7.**
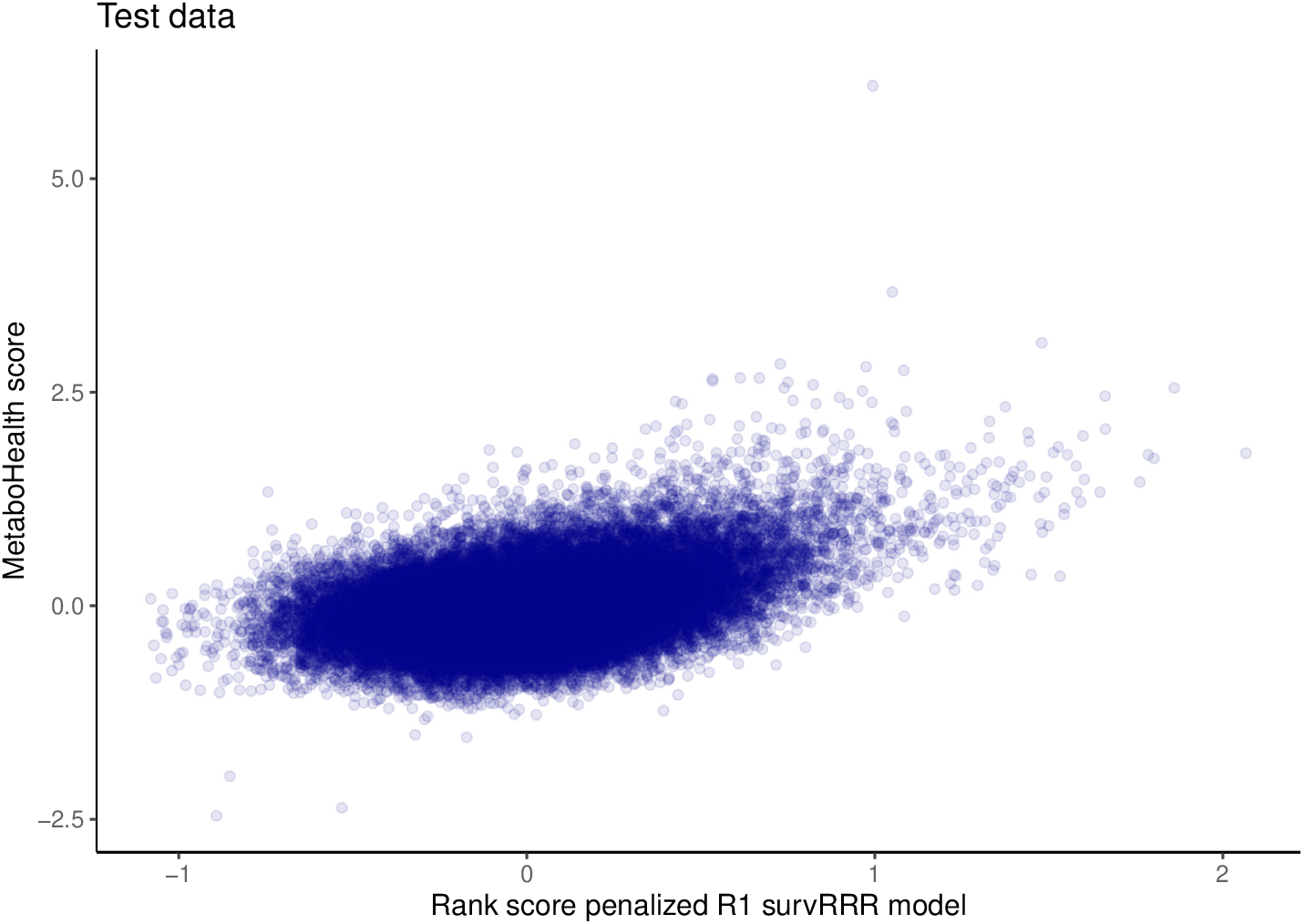
Comparison of rank scores from optimal penalized survRRR model, which is of rank 1, with the MetaboHealth score. Both models were fitted on the same training data set. Opacity of the dots was set to 0.1.

## 4 Discussion

In this paper we introduced a penalized reduced rank regression model to analyse multi-outcome survival data. We call this model the penalized survRRR model. By constraining the coefficient matrix 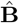 to be of max rank *R*, the reduced rank model assumes that there is a set of *R* latent factors that drive all outcomes considered. We introduced a penalized version of this model to allow for the inclusion of high-dimensional and/or strongly correlated predictors or outcomes. Depending on the problem at hand, different penalties can be considered. We discussed the ridge- and lasso-penalized model in detail and provided proof that the lasso-penalized survRRR model results in unique **A** and **Γ** matrices, in contrast to the unpenalized or ridge-penalized survRRR model.

The penalized survRRR model is a valuable addition to the methodological toolkit of approaches that can be used to analyze multimorbidity data (or, in fact, any type of multi-outcome survival data). As multimorbidity is becoming the rule rather than the exception^27^, there is an increasing interest in discovering and gaining insight in the shared underlying biological mechanisms of age-related diseases and death. This is exactly what the penalized survRRR model can provide. In the context of ageing, an appealing characteristic of the penalized survRRR model is that the optimal number of ranks can be deduced from the model. This provides insight into the multi-faceted nature of the human ageing process. If the optimal number of ranks of such a model is equal to 1, it provides support to the convention of grouping all considered diseases under the ‘age-related’ denominator, and to the current practice of only considering time-to-death when developing a predictor of ageing. If the optimal number of ranks of a penalized survRRR model fitted on multiple time-to-age-related diseases or death is greater than 1, it can be argued that a model built on time-to-death alone will miss relevant aspects of ageing.

We fitted the lasso-penalized survRRR model on multimorbidity data from the UK Biobank, using as predictors 242 blood-based metabolic variables and as outcomes time to seven different age-related diseases and death. We found that the rank 1 model best describes the data. The optimal model assigned a weight of 0 to seven of the 242 metabolic variables and to none of the eight outcomes. Concretely, this means that a single latent factor drives all eight outcomes considered. This underpins the conjecture that all considered diseases can indeed be grouped in the same category of ‘age-related’ diseases, when using the considered metabolic variables as predictors. The results also provide support to the current practice within the ageing field to develop (biological) age predictors based on time-to-mortality alone. Our results indicate that this is a reasonable approach when using metabolic variables as predictors, as no second latent factor was found that drives some of the age-related diseases we considered but not death. In such a scenario, that second latent factor would be missed when fitting a model on time-to-mortality only. However, we also illustrated that the rank scores of our lasso-penalized survRRR model captured more than a model fitted on time-to-mortality alone.

Even though we found that a rank 1 model best describes the data we considered, our results do not contradict the fact that ageing is an considered an inherently multifaceted phenomenon^4^. If reality is actually of a higher rank, but one rank dominates (i.e. the eigenvalues of the other ranks are very low), the rank 1 model might provide the best, most stable fit. This could be the case in our motivating application as well. Additionally, we only considered a limited set of metabolic variables as predictors, and just seven age-related diseases. Ageing researchers might therefore be interested in fitting the penalized survRRR model to other omics data, or even multiple omics simultaneously, and in extending the number of age-related diseases considered.

In this work we considered the two most well-known penalties: ridge and lasso. However, in principle the survRRR model can be combined with any type of penalty. The extension to other penalty types, e.g. (sparse) group lasso, is left as future research. This requires some additional work: for this current implementation, we could make use of existing functions in the statistical programming language R to fit a stratified lasso-penalized Cox proportional hazards model. To the best of our knowledge, there does not yet exist an implementation of a stratified group lasso-penalized Cox model. We are currently investigating which algorithmic approaches would be most suitable for this task.

## Acknowledgements

We thank Myriana Miltiadous for checking the penalized survRRR R-function and her contributions to the code for the simulated example data set.

## Data and code availability statement

UK Biobank data are publicly available to bona fide researchers upon application at http://www.ukbiobank.ac.uk/using-the-resource/. Software for the penalized survRRR model in the form of R code, together with a simulated example data set, is available at https://github.com/marije-sluiskes/penalized_survRRR and is provided as Supporting Information.

