## Supplementary material for "Penalized reduced rank regression for multi-outcome survival data supports a common metabolic risk score for age-related diseases": SuppInfo_Appendix_A.pdf

The calculation of the MetaboHealth score as described below follows the procedure outlined in the original publication by Deelen *and others* (2019):

1. Use the same training and test data sets as defined when fitting the penalized survRRR model. As we do not refit the MetaboHealth model, we only consider the test data set.

2. In the test data set, start from the original (raw) metabolic variable values. (For MetaboHealth, the metabolic variables need to be log-transformed before centering and scaling. When fitting the penalized survRRR model, metabolic variables were centered and scaled without a log-transformation.)

3. Add a value of 1 to the raw metabolic variables, log-transform, center and scale.

4. Calculate the MetaboHealth score as: $\mathit{MetaboHealth} = \log(0.8) \times \mathit{xxl\_vldl\_l} + \log(0.87) \times \mathit{s\_hdl\_l} + \log(0.85) \times \mathit{vldl\_size} + \log(0.78) \times \mathit{pufa\_pct} + \log(1.16) \times \mathit{glucose} + \log(1.06) \times \mathit{lactate} + \log(0.93) \times \mathit{his} + \log(1.23) \times \mathit{ile} + \log(0.82) \times \mathit{leu} + \log(0.87) \times \mathit{val} + \log(1.13) \times \mathit{phe} + \log(1.08) \times \mathit{acetoacetate} + \log(0.89) \times \mathit{albumin} + \log(1.32) \times \mathit{glyca}$. Here, the metabolite names already refer to the transformed metabolites as defined under step 3.
